# Stromal signals dominate gene expression signature scores that aim to describe cancer-intrinsic stemness or mesenchymality characteristics

**DOI:** 10.1101/2023.08.25.554747

**Authors:** Julian Kreis, Bogac Aybey, Felix Geist, Benedikt Brors, Eike Staub

## Abstract

**Purpose:** Epithelial-to-mesenchymal transition (EMT) in cancer cells confers migratory ability, a crucial aspect of tumor metastasis that frequently leads to death. In multiple studies, authors proposed gene expression signatures for EMT, stemness, and mesenchymality (EMT-related) characteristics of tumors based on bulk tumor expression profiling. However, recent studies have suggested that non-cancerous cells in the tumor micro- or macroenvironment heavily influence individual signature profiles.

**Experimental Design:** We analyzed scores of 11 published and frequently referenced gene expression signatures in bulk, single cell, and pseudo bulk expression data across multiple cancer types.

**Results:** Our study strengthens and extends the influence of non-cancerous cells on signatures that were proposed to describe EMT-related (EMT, mesenchymal, or stemness) characteristics in various cancer types. The cell type composition, especially the amount of tumor cells, of a tumor sample frequently dominates EMT-related signature scores. Additionally, our analyses revealed that stromal cells, most often fibroblasts, are the main drivers of the EMT-related signature scores.

**Conclusions:** We call attention to the risk of false conclusions about tumor properties when interpreting EMT-related signatures, especially in a clinical setting: high patient scores of EMT-related signatures or calls of “stemness subtypes” often result from low tumor cell content in tumor biopsies rather than cancer cell-specific stemness or mesenchymality/EMT characteristics.

## 1. Introduction

The ability to switch from a stationary epithelial cell state to a motile mesenchymal cell state, called epithelial-to-mesenchymal transition (EMT), is essential for stem cell differentiation and de-differentiation. This capability is a key driver of tumor cell plasticity for tumor cells, which is required for tumor cells to invade distant tissues and metastasize [1]. Using bulk RNA-seq data, multiple gene expression signatures have been postulated to describe stemness [2, 3], mesenchymality [4–7], and EMT [8–12] properties (hereafter referred to as EMT-related signatures) of cancer cells in colorectal (CRC), breast (BRCA), glioblastoma (GBM), and head and neck cancers (HNC). Subsequently, pioneering cancer characterization programs from The Cancer Genome Atlas (TCGA) [13] have applied these signatures for cancer subtype identification or characterization of cancer-specific traits.

However, the expression profiles of bulk RNA-seq samples represent a complex mixture of pathway activity within cells in combination with heterogeneous cell-type compositions. In samples with a high tumor content, expression signatures may provide high precision for tumor-intrinsic molecular states. In samples with comparably low tumor content, differences in cell composition between samples could dominate the scores of signatures originally intended to measure tumor-intrinsic properties. Multiple studies have highlighted the strong dependency of selected gene expression signatures on tumor microenvironment (TME) composition, limiting their applicability for cancer subtype stratification [14, 15], clinical study designs [16–18], and preclinical cancer models [12]. Similarly, tumor and stromal cell expression profiles from microdissected and single-cell RNA-seq (scRNA-seq) samples from CRC [19–22], ovarian [23], HNC [14], and pan-cancer [24] studies reemphasized such dependencies and identified fibroblasts as crucial contributors to the elevated scores of different EMT-related signatures, which were used to stratify patients into mesenchymal subtypes. Some research groups have reinterpreted the influence of fibroblast content on mesenchymal subtypes as a sign of fibrosis [20] and others as artifacts in the sampling procedure [14]. Eide et al. argued that cancer cell-intrinsic processes control stromal composition and maintain the biased subtype by partly correcting for stromal contamination in CRC tumors [12]. This conceptual problem is evident for several cancer indications; however, consensus on the experimental procedures required for capturing EMT-related tumor subtypes is lacking. In addition, a comprehensive analysis of a large panel of EMT/mesenchymality-related subtype signatures in the presence of varying tumor content in samples is lacking.

Here, we dissected the influence of cells in the surrounding tumor tissue (i.e., the micro- and macroenvironment) with a focus on the influence of fibroblasts and immune cells on commonly used gene expression signatures for EMT-related phenotypes. To this end, we used RNA-seq data from both bulk and single cells. Using RosettaSX, our published platform for gene expression signature scoring, and comprehensive analyses of scRNA-seq data, we investigated the association between EMT-related signature profiles and tumor/stroma content in samples of different tumor indications. We assessed the risk of premature conclusions when not considering the cell type composition of tumor samples in the assessment of EMT-related gene expression signatures, specifically for HNC, BRCA, GBM, and CRC.

## 2. Results

Previous studies have shown that the expression profiles of individual EMT-related signatures across clinical tumors are highly congruent but could be strongly influenced by non-cancerous cells in the TME or simply by tumor sample composition (tumor macroenvironment, normal stromal tissue surrounding a tumor). We used the workflow shown in Figure 1 to analyze 11 different EMT, mesenchymal, and stemness gene expression signatures (hereafter referred to as EMT signatures) in bulk tumors, cell lines, and single-cell gene expression data to identify the factors that primarily drive their scores [2–12].

**Figure 1:**
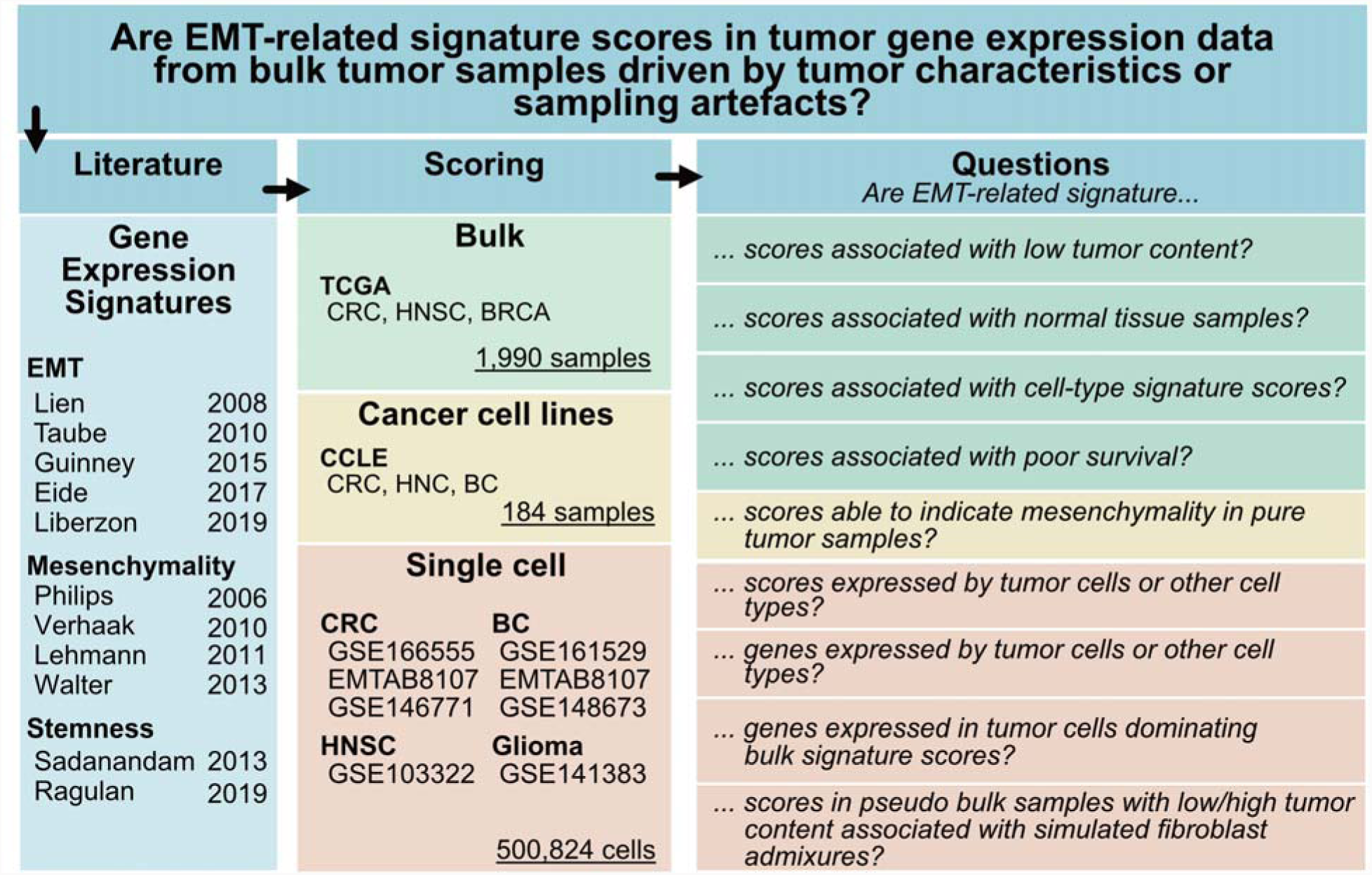
Workflow for delineating the influence of non-tumor cells on EMT-related signature scores. We identified 11 frequently applied gene expression signatures from literature that were proposed as markers for stemness or mesenchymality character of tumors. We investigated the signatures in gene expression data of clinical samples, cancer cell lines, and single cells to address questions related to their utility for clinical cancer profiling. EMT: epithelial-to-mesenchymal transition, TCGA: the cancer genome atlas, CCLE: Cancer cell line Encyclopedia

### 2.1 Diverse EMT, mesenchymal, or stemness signatures generate highly congruent signature profiles in primary cancer patients

First, we assessed the association of 11 EMT signature profiles with tumor content across samples and analyzed them in the context of a literature-derived collection of gene expression signatures in our RosettaSX system [25]. We focused on the analysis of the CRC, BRCA, and HNC bulk RNA-seq data. Our signature collection enabled us to explore the associations between EMT signatures and other signature profiles for a variety of cancer-relevant processes. Next, using normal tissue adjacent to the tumor (NAT), cancer cell line bulk RNA-seq, single-cell RNA-seq (scRNA-seq), and simulated pseudo-bulk data, we investigated the contribution of non-cancerous cells from the macro- or microenvironment of tumors to EMT-related signature scores.

To assess the signature landscape of CRC, we analyzed 380 TCGA CRC tumors (TCGA COAD and READ) using RosettaSX. We applied the coherence score (CS) [25, 26] to determine the within-signature congruence of gene expression profiles. The CS indicates whether genes in a gene expression module are coordinately activated across patients. This indicates whether a given signature is coherently active in some samples while being inactive in others. Only signatures with high CSs should be used to infer the activity of their underlying biological phenomena in a single sample of the dataset to be explored. A total of 121 of the 285 RosettaSX signatures were coherently expressed (CS > 0.18, at least 50 % of genes available). From these, we selected signatures relevant to known cancer-related processes (e.g., proliferation, and CRC subtypes), cellular composition of the tumor sample, and EMT- or stemness-related signatures. Furthermore, we filtered out signatures with a Jaccard index larger than 0.25 to other signatures to remove overly redundant biological processes. Thereby, we derived a set of 44 signatures that comprise the 11 published EMT signatures and cover other biological phenomena, such as the expression of extracellular matrix (ECM) or proliferation genes, or the presence of specific cell-types, such as T-, B-cells, or fibroblasts (Figure 2, left side).

**Figure 2:**
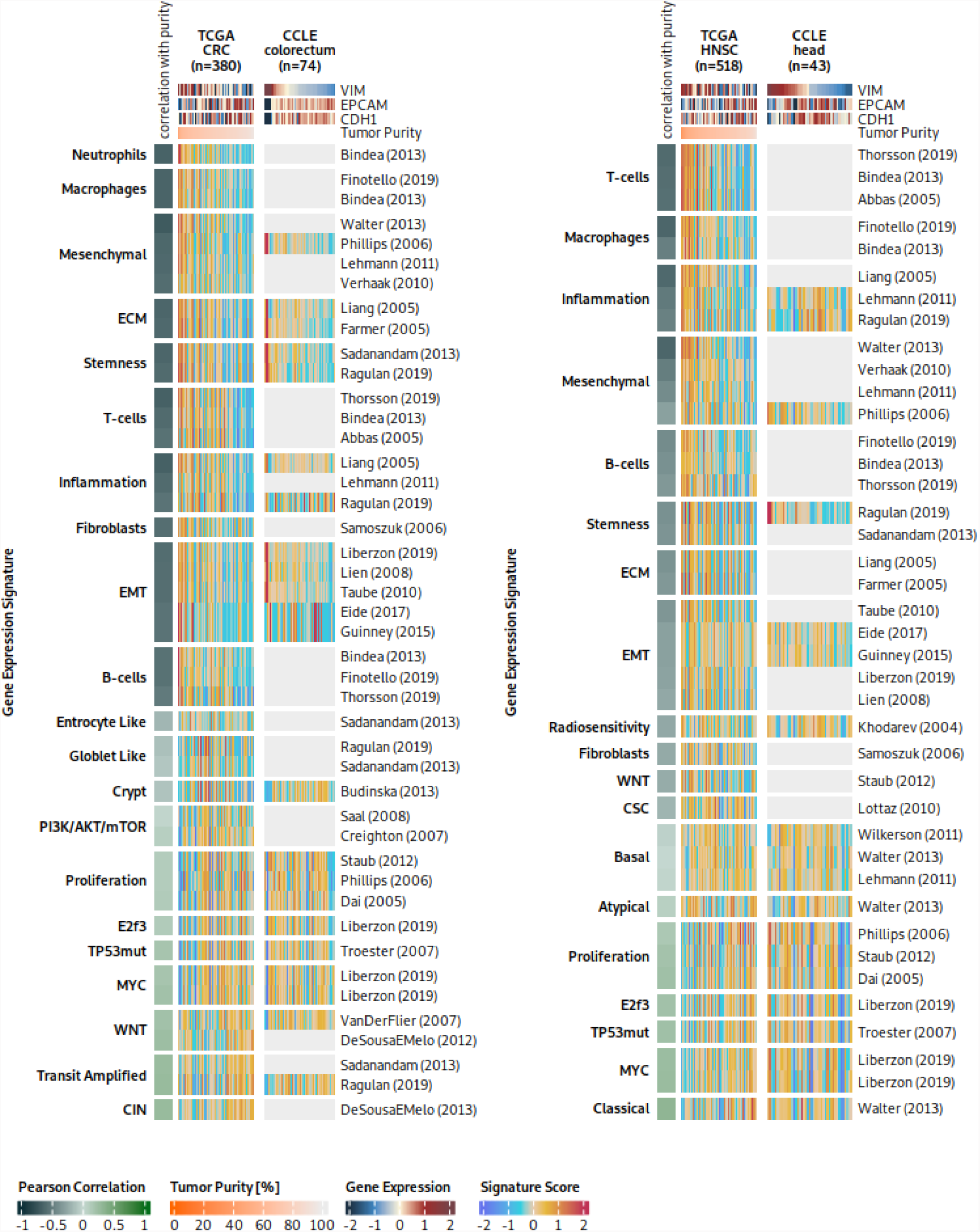
EMT-related signatures and their associations with cancer pathways or processes. We studied 11 mesenchymal, EMT, and stemness signatures and their relation to other biological processes in TCGA cancer patients and CCLE cancer cell lines. Left heatmaps: Gene expression signature scores of coherent signatures and their correlation with sample purity (CPE) in TCGA CRC samples. The heatmaps have been supplemented with single gene marker expression. The column annotation lists important markers for mesenchymal (VIM) and epithelial (EPCAM, CDH1) properties. Samples from left to right have decreasing levels of vimentin expression, and signatures groups have an increasing correlation with tumor purity from bottom to top. Light grey areas indicate signatures that did not satisfy the coherence scoring threshold. Right heatmaps: An analogous analysis as in CRC to the left, here for TCGA HNC samples.

The CS analysis revealed that the 11 EMT-related signatures had a median (MED) CS of 0.41 (interquartile-range (IQR) = 0.26), whereas the remaining 33 signatures reached a median CS of 0.32 (IQR = 0.22). Thus, genes of EMT-related signatures are among the signatures with the strongest gene co-expression, indicating a coordinately regulated expression pattern that is presumably guided by a strong underlying biological phenomenon involving EMT signature genes. EMT-related signature scores were more strongly correlated with common mesenchymal markers than with epithelial markers (one-sided Wilcoxon test, p<.0001, effect size = 0.85), suggesting a predominantly mesenchymal characteristic of mesenchymal-related signatures as well as stemness and EMT signatures. In the CRC bulk RNA-seq data, EMT-related signature profiles were strongly associated with signatures describing the presence of different non-cancerous cell types (macrophages [27], fibroblasts [28], or B-cells [29]) and genes encoding the ECM [30, 31]. These associations might indicate that cells in the tumor environment (microenvironment or tumor-adjacent normal macroenvironment) strongly contribute to EMT-related signature scores. We observed an anti-correlation between gene expression signatures and phenomena such as MYC activation [11], proliferation [26, 32], and CRC subtypes or cell-of-origin (e.g., goblet-like, crypt, or CIN) gene expression signatures [2, 3, 33].

In the TCGA HNC gene expression data, 40 signatures met the previously described selection criteria. Their signature scores recapitulated our findings regarding CRC (Figure 2, right panel). EMT-related signatures were more strongly correlated with non-cancer cell-type signatures (e.g., T-cell and B-cell) than with cancer-associated processes (e.g., proliferation, and MYC activation). Similarly, EMT-related signatures were more strongly correlated with mesenchymal markers than with epithelial markers (one-sided Wilcoxon test, p<.0001, effect size 0.85). Analogous analyses of TCGA BRCA data yielded comparable results (Figure S1).

In summary, these results indicate that non-cancerous cells in the tumor macro- or microenvironment contribute strongly and sometimes even dominantly to EMT-related signature scores. Therefore, we decided to further investigate whether the signals of EMT signatures are generally associated with tumor sample impurities, that is, the low tumor content of the samples.

### 2.2 EMT-related signatures scores are high in samples with low tumor content

Bulk RNA-seq tumor samples can comprise high fractions of tumor-adjacent normal tissue cells, especially stromal and immune cells, if no macro- or microdissection is performed by pathologists during sample preparation. Stromal or immune cells of tumor-adjacent tissues can potentially influence gene expression signals more strongly than the characteristics of cancer cells themselves. Thus, we compared TCGA-provided tumor content with signature scores in the TCGA CRC population (Figure 2 left, Figure S2 A, B). EMT and immune cell type signature scores were strongly anti-correlated with tumor content (rMED(EMT) = -0.6 and rMED(IMM)=-0.62), whereas the other signatures were weakly positively correlated with tumor content (rMED = 0.11). A comparison of the 11 EMT signatures with the 22 cancer process signatures revealed that EMT signatures had a stronger negative association with tumor content than any other process (excluding cell type signatures, one-sided Wilcoxon test, p=1e-04, effect size = 0.67). Additionally, there was a strong correlation between the signature scores of EMT-related signatures and the stromal signature for BRCA (R between 0.7 to 0.98; signature from Farmer et al. 2005). These findings suggest that the presence of non-tumor cells from the stroma significantly drives, if not dominates, EMT-related signature scores. Analyses of other cancer types represented in TCGA yielded comparable results (Figure 2, right panel, and Figure S1).

Summarizing our results on the bulk gene expression data of cancer patients, we found that the collection of 11 EMT- or stemness-related signatures (a) each had a stable expression footprint (high coherence score) in bulk gene expression data, (b) shared very similar signature profiles in clinical tumor specimens despite differences in the procedures for their discovery and their gene sets, (c) were significantly associated with low tumor content, and (d) all correlated strongly with a well-known signature to track stromal content in BRCA. These findings motivated us to investigate 11 EMT signatures in pure tumor cells (cancer cell lines) and single cell gene expression datasets.

### 2.3 EMT, mesenchymal, and stemness signatures are less congruent in TME-naïve cancer cell line models than in bulk samples

To investigate whether TME-naïve pure cancer models (lacking immune or stromal cells) show signs of differential EMT states or mesenchymality/stemness, we analyzed our set of EMT-related gene expression signatures in CRC cell line data (Figure 2 left, right heatmap). Of the 44 signatures we previously selected for CRC TCGA samples, only 22 passed our quality control procedure based on the coherence score. As expected, many cell type gene sets (e.g., T-cells, B-cells, and macrophages) have a low coherence score, and consequently, interpretations of signals of these signatures in the cell line data cannot be regarded as meaningful.

However, EMT-related signatures yielded acceptable coherence scores. Our analysis revealed a small group of cell lines with extraordinarily high signature scores for the 11 EMT-related signatures and signatures related to the ECM or fibroblasts (Figure 2 B, Figure S2 C). These cell lines exhibited low epithelial marker expression (EPCAM) and high mesenchymal marker expression (CDH1). Their results warrant follow-up to investigate whether EMT-high cell lines are of epithelial origin, and a review of their annotations revealed that three cell lines originated from mesenchymal cells but not carcinoma cells (HS255T, HS675T, and HS698T; from fibroblasts or from a gastric tumor, presumably a gastrointestinal stromal tumor). Removal of these cell lines and restriction of the analysis to carcinoma cell lines drastically worsened the coherence scores, with an average CS decrease of 0.11 in the EMT signatures (Figure S3). This indicated that the higher coherence scores in the data for all 44 cell lines were primarily driven by cell lines that originated from mesenchymal cancers, not colorectal carcinoma. This not only exemplifies the utility of the coherence score approach, but also further questions whether the investigated EMT signatures capture expression signals of mesenchymality that are intrinsic to the tumor.

### 2.4 Tumor macroenvironment can influence EMT-related signature scores

Next, we compared normal tissue adjacent to the tumor (NAT) with samples with high and low tumor content to study the influence of the tumor macroenvironment (Figure 3). As shown before, it was evident that across cancer indications, EMT-related signature scores were most elevated in samples with low tumor content. If the tumor content was high, the scores were among the lowest across all the samples.

**Figure 3:**
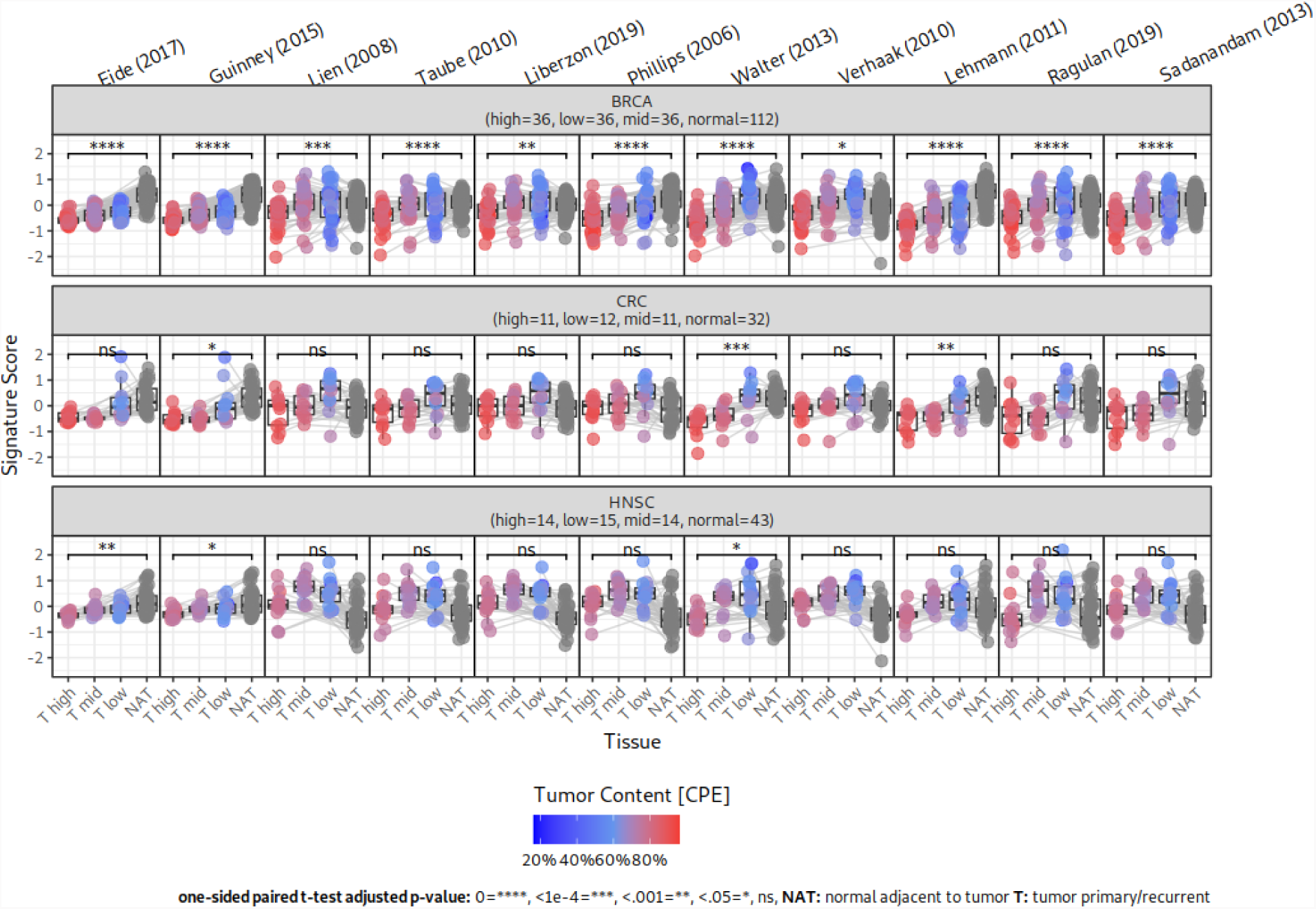
Comparison of EMT-related gene expression signature scores in tumor and normal tissue that is adjacent to the tumor tissue. Tumor samples are separated into low, medium (med) and high tumor content (CPE) based on the 25th and 75th percentiles of TCGA tumor content estimates. Lines indicate the paired tumor and normal adjacent to tumor (NAT) samples. Colors indicate the tumor content in tumor samples with low (blue), or high (red) tumor content or grey for missing data. Scores in NAT and high tumor content samples were compared using paired t-tests. P-values were adjusted using Bonferroni correction.

In BRCA, all EMT-related signature scores were significantly higher in NAT than in samples with high tumor content (Figure 3, first row), but this trend was not observed in CRC and HNSC. The only exceptions were the signatures reported by Walter et al. and Lehman et al.. These two signatures had significantly higher scores in the NAT within high tumor content samples for at least two indications. In HNSC, almost all analyzed signatures had low scores in NAT tissue. This might be due to the overall low tumor content in the analyzed HNSC samples.

These results highlight that within individual indications, signatures might not only be guided by cells in the tumor microenvironment, but also by the macroenvironment (concurrently sampled normal tissue), especially in BRCA and CRC.

### 2.5 Cell type composition strongly affects scores of EMT-related signatures

To further investigate the origin of elevated EMT-related signature scores in bulk tumor specimens, we analyzed their signature scores and individual signature gene expression levels in eight scRNA-seq datasets derived from CRC, BRCA, glioblastoma, and Head and Neck squamous cell carcinoma (HNSC) samples (see Methods). After analyzing the impact of the macroenvironment on EMT-related signature scores in the previous sections, we analyzed the impact of cell type-specific expression profiles in the TME.

First, we examined the expression of EMT-related signature scores within the individual cell types (Figure 4). With high consistency across the four analyzed cancer indications, we found that all EMT-related signatures displayed the highest average expression in cell populations of the TME, fibroblasts, myofibroblasts, or endothelial cells, but not in malignant cells. These high expression levels in fibroblasts underpin the overall dependency of the EMT signatures on stromal cell signals but not on cancer cells. Overall, only a fraction of the cancer cells expressed EMT-related signatures at relatively low levels.

**Figure 4:**
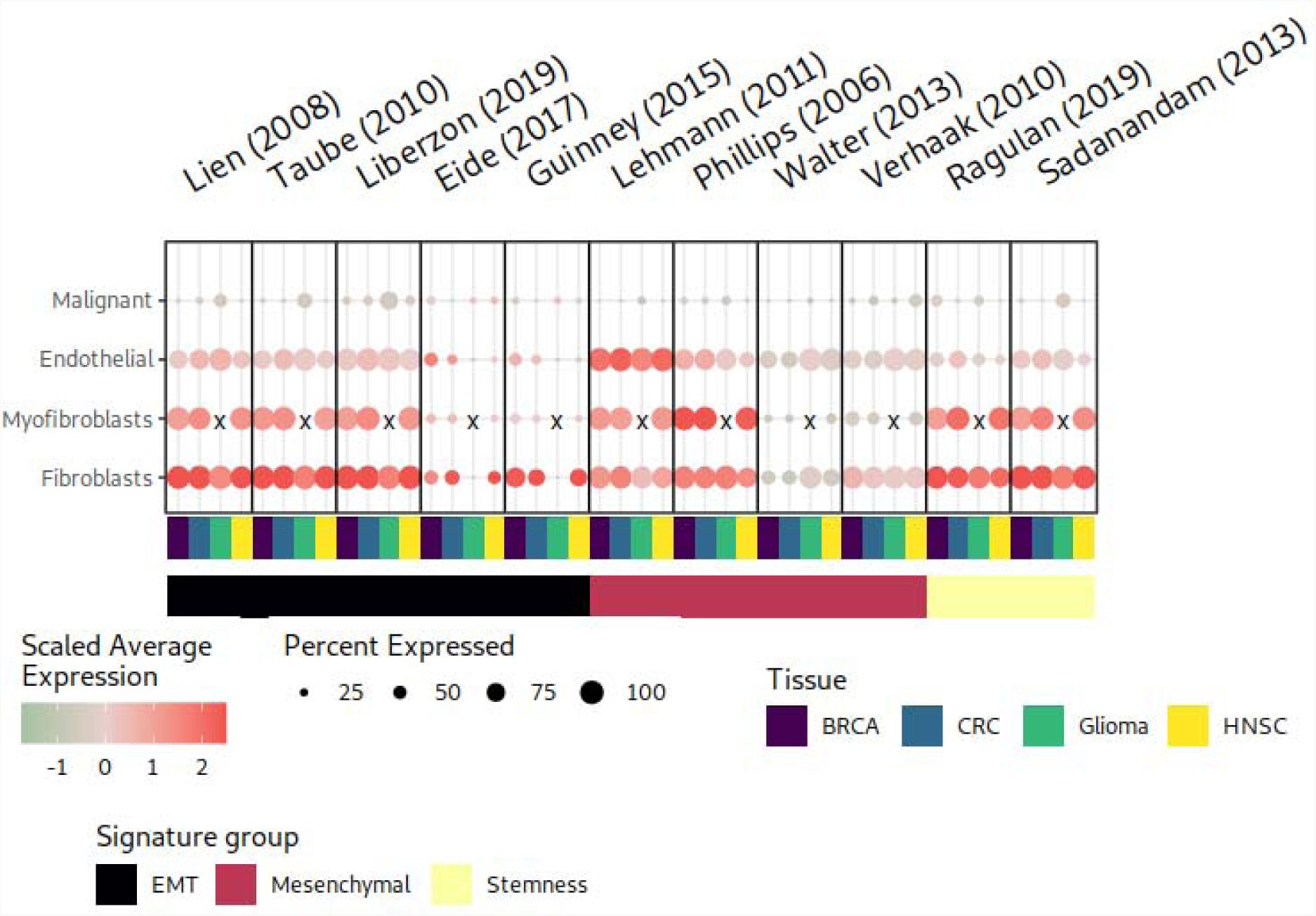
Comparison of EMT-related signature scores in eight scRNA-seq datasets. The dot plot shows the module scores of EMT-related signatures and the number of cells expressing them in the different cell types. Each panel displays dot plots to quantify the expression of individual signatures in four different tissues (malignant, endothelial, myofibroblast, and fibroblast) derived from scRNA-seq datasets for BRCA, CRC, glioma, and HNSC. x: Unavailable cell type

### 2.6 High EMT-related signature scores are associated with fibroblast enriched pseudo bulk samples

If tissue biopsies are used in combination with bulk sequencing experiments, we hypothesized that minor sampling artifacts can strongly influence signature scores depending on the fraction of sampled fibroblasts. To further study the dependency of EMT-related signature scores on the fraction of fibroblast cells, we generated 20 pseudo-bulk RNA-seq samples based on CRC scRNA-seq data from Lee et al.. Each pseudo bulk sample was composed of RNA counts from different ratios of tumor to fibroblast cells, ranging from 20% to 80% of tumor content (see Methods). We investigated the similarities of RosettaSX pathway and process signatures in an integrated analysis of 20 pseudo-bulk and 380 TCGA CRC bulk samples using an integrated RosettaSX analysis (see Methods, Figure 5). Pseudo-bulk samples with enriched fibroblast content co-clustered with samples with low tumor content They had elevated EMT-related, stroma, or TME cell type signature scores, indicating that fibroblasts might significantly drive the signature scores of the analyzed bulk RNA-seq samples and drive co-clustering. In contrast, bulk samples with high tumor content had elevated scores for proliferation or CRC subtype signatures (e.g., transit amplified [3], and crypt [34]). Our results showed that high EMT-related signature scores in low-purity TCGA samples can result from abundant fibroblast cells in the low-tumor-content samples and do not necessarily originate from the mesenchymality of the cancer cells.

**Figure 5:**
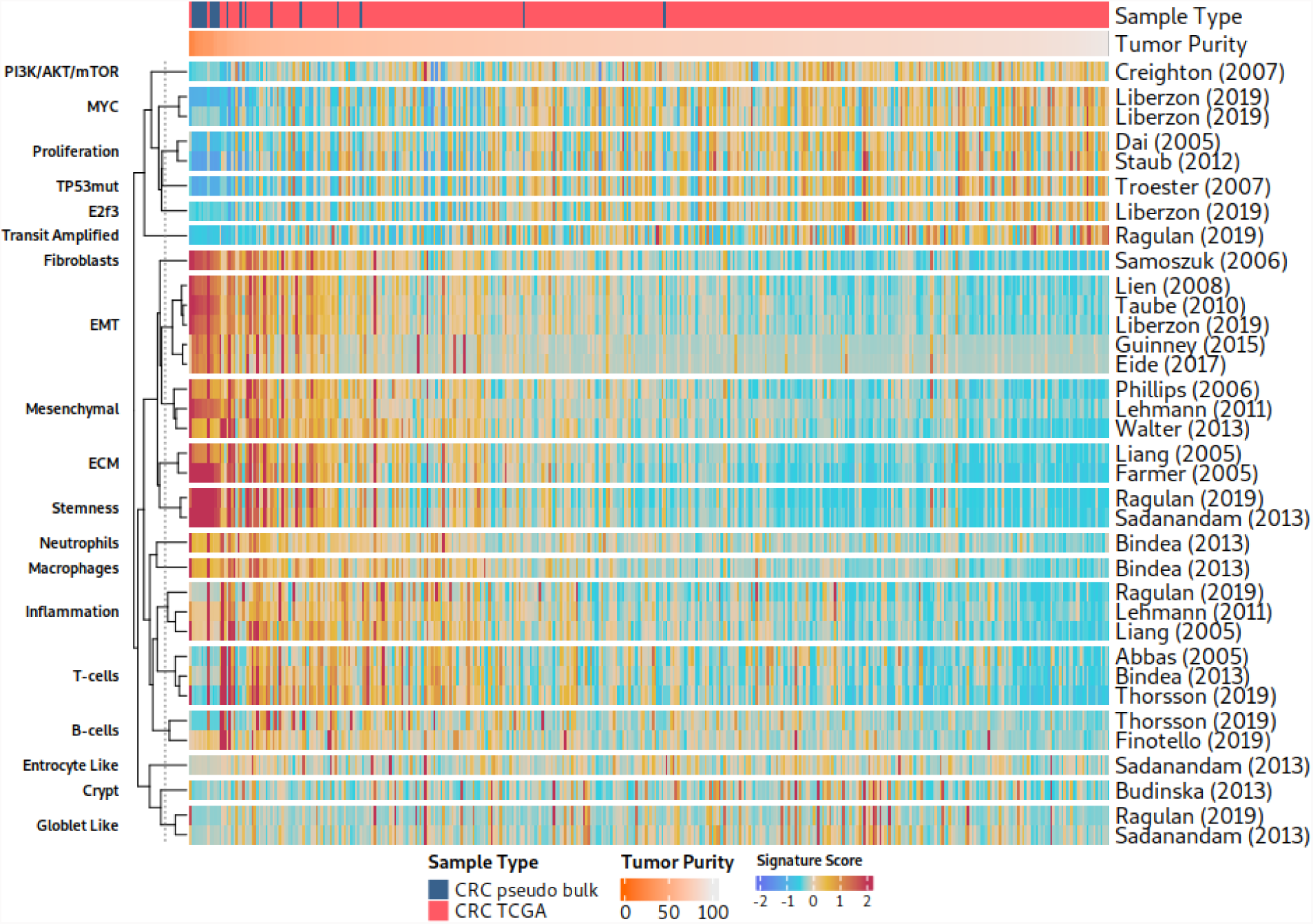
RosettaSX analysis of an integrated dataset of TCGA CRC and fibroblast enriched pseudo bulk samples. Top annotation indicates the tumor purity (CPE) for TCGA samples, and the fraction of fibroblast cell RNA used for pseudo bulk generation (see methods). The tumor samples (columns) are sorted by their tumor purity from low (left) to high (right). Note the tight clustering of all 11 EMT-related signatures and their correlation with tumor purity.)

### 2.7 Fibroblasts dominate the expression for most genes in EMT-related signatures

Furthermore, we investigated the cell-type specificity of the individual signature genes(Table 1). Overall, the results of our investigations of cell-type-specific expression of the signature genes were congruent across the analyzed CRC, breast, HNC, and glioma scRNA-seq datasets. Most of the cell types that predominantly expressed individual signature genes were fibroblasts, but not tumor cells (Figure S4). Differential gene expression analysis of all available signature genes within the eight scRNA-seq datasets confirmed the low contribution of tumor cells to the signature scores (see Table 1). Overall, only a small number of genes showed significantly higher expression in cancer cells than in fibroblasts. However a much higher fraction of genes showed significantly higher expression in fibroblasts. Only four signatures for mesenchymal and EMT characteristics indicated a higher expression of a few genes (a maximum of 3.7 % of individual signature genes) in cancer than in other cell types in at least one dataset [4, 5, 8, 11]. These patterns highlight that fibroblasts were the primary contributors to the high overall signature scores for our set of EMT-related signatures. Thus, these EMT-related signatures often predominantly describe mesenchymal characteristics of the tumor macro- or microenvironments rather than the stemness or mesenchymal character of cancer cells when bulk tumor tissue is profiled, especially when the tumor content is low and the fraction of adjacent normal tissue is high.

**Table 1:** Differentially expressed EMT-related signature genes between fibroblasts and cancer cells in summarized across three (BRCA), three (CRC), one (glioma), and one (HNC) scRNA-seq datasets.

| **Geneset** | **BRCA<sup>1</sup>** | **CRC<sup>1</sup>** | **Glioma<sup>1</sup>** | **HNSC<sup>1</sup>** |
| --- | --- | --- | --- | --- |
| **EMT** |  |  |  |  |
| Eide (2017) | 2.1 (0.0) | 2.6 (0.0) | 0.0 (0.0) | 5.6 (0.0) |
| Guinney (2015) | 6.7 (0.0) | 8.3 (0.0) | 0.0 (0.0) | 10.5 (0.0) |
| Liberzon (2019) | 25.5 (0.2) | 29.0 (0.0) | 13.2 (0.5) | 18.4 (1.0) |
| Lien (2008) | 38.3 (0.0) | 45.7 (0.0) | 14.8 (0.0) | 33.3 (0.0) |
| Taube (2010) | 25.2 (0.4) | 26.6 (0.0) | 17.3 (0.0) | 18.2 (1.1) |
| **Mesenchymal** |  |  |  |  |
| Lehmann (2011) | 3.4 (0.0) | 5.7 (0.0) | 2.9 (0.0) | 3.5 (0.0) |
| Phillips (2006) | 56.4 (0.0) | 64.1 (0.0) | 38.5 (0.0) | 38.5 (0.0) |
| **EMT** |  |  |  |  |
| Verhaak (2010) | 3.7 (0.0) | 4.1 (0.0) | 2.6 (0.0) | 2.6 (0.7) |
| Walter (2013) | 5.6 (0.3) | 6.3 (0.0) | 2.8 (0.0) | 5.2 (0.4) |
| **Stemness** |  |  |  |  |
| Ragulan (2019) | 44.4 (3.7) | 53.7 (0.0) | 25.0 (0.0) | 44.4 (0.0) |
| Sadanandam (2013) | 27.4 (0.5) | 35.7 (0.0) | 8.2 (0.5) | 25.9 (0.0) |

<sup>1</sup> DEG Fibroblast % (DEG Malignant cells %)

### 2.8 Cancer cell-specific gene expression does not dominate bulk sequencing signature scores of EMT-related signatures

Our single-cell analysis showed a high expression of most EMT-related signature genes in the TME cells. Thus, it is likely that the signature score profiles in bulk sequencing data are primarily driven by TME cells. To translate the findings of our single-cell analysis to bulk sequencing experiments, for each signature gene, we compared three variables: 1) expression fold changes between tumor and fibroblast cells of scRNA-seq data, 2) the correlation of the gene expression profile with tumor purity (bulk RNA-seq data), and 3) the correlation of the gene expression profile with the respective signature scores in TCGA CRC patients (Figure 6).

**Figure 6:**
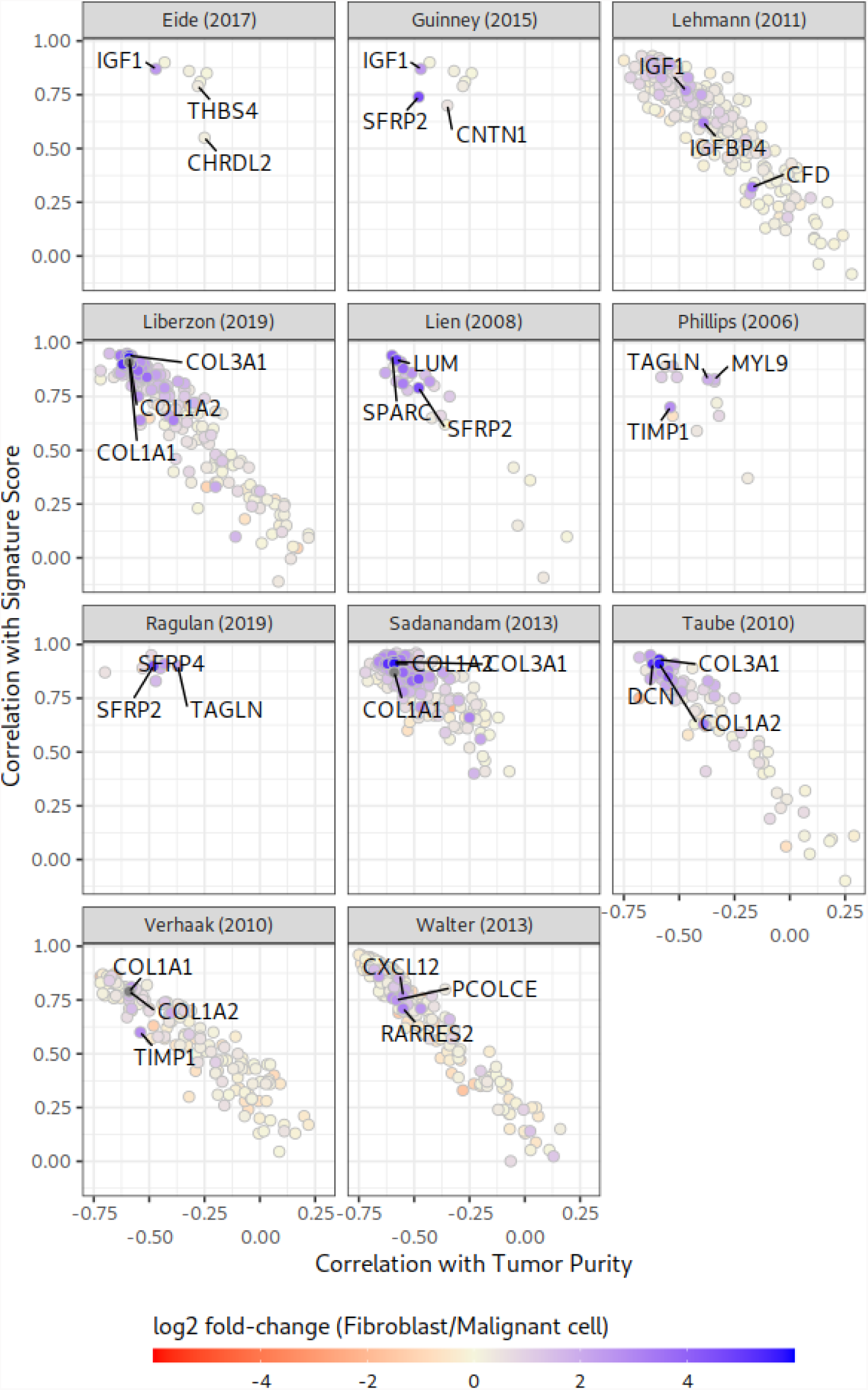
Influence and association of signature gene expression profiles on signature scores in CRC bulk RNA-seq samples. Each dot in a panel represents a signature gene of the respective signature. A gene at the bottom weakly influences the overall signature score. A gene at the top strongly influences the overall signature score. Genes on the left are stronger correlated with low tumor content. Blue coloring of data points indicates a higher expression in fibroblasts than in malignant cells in our scRNA-seq analyses.

Overall, genes strongly associated with the respective signature scores (in bulk sequencing data) were expressed at higher levels in fibroblasts than in tumor cells in the scRNA-seq analysis. Additionally, at the bulk sequencing level, genes with high association with the respective signature score and high fibroblast expression were associated with low tumor purity. These patterns indicate that genes associated with low tumor purity significantly contribute to EMT-related signature scores, and that fibroblasts express many of these genes at a higher level than tumor cells (Figure 6). The genes that were highly expressed in fibroblasts were VIM, COL1A1, COL1A2, and LUM, which have been previously associated with fibroblasts [35, 36].

### 2.9 Our analyses do not confirm a prognostic value of EMT-related gene expression signature scores

Previous studies of individual EMT-related signatures have indicated an association between high scores and poor survival [2, 19]. Here, we re-evaluated the association of signature scores with survival signals, specifically with disease-free interval (DFI) survival, in CRC and BRCA TCGA patients (Table 2, Table S1). We modeled the influences of the signature scores on DFI using univariate and multivariate Cox proportional hazard models, accounting for tumor stage. Neither the univariate nor the multivariate analysis indicated significant associations. Similarly, there was no significant association between survival and tumor content.

**Table 2:** Cox proportional hazards analysis for EMT-related signature scores and TCGA cancer patients. We show the results of univariate and multivariate Cox regression models to analyze the influence of EMT-related gene expression signature scores and stage annotation on progression-free interval survival of TCGA CRC patients. Multivariate analysis was performed using tumor stage as a covariate.

| COAD |  |  |  |  |  |  |
| --- | --- | --- | --- | --- | --- | --- |
| Characteristic | Univariate |  |  | Multivariate |  |  |
|  | HR<sup>1</sup> | 95% CI<sup>1</sup> | p-value | HR<sup>1</sup> | 95% CI<sup>1</sup> | p-value |
| **EMT** |  |  |  |  |  |  |
| Lien (2008) | 0.76 | 0.32, 1.78 | 0.5 | 0.68 | 0.27, 1.72 | 0.4 |
| Taube (2010) | 0.72 | 0.29, 1.80 | 0.5 | 0.64 | 0.24, 1.73 | 0.4 |
| Liberzon (2019) | 0.75 | 0.29, 1.92 | 0.5 | 0.66 | 0.24, 1.84 | 0.4 |

COAD
| Characteristic | Univariate |  |  | Multivariate |  |  |
| --- | --- | --- | --- | --- | --- | --- |
|  | HR<sup>1</sup> | 95% CI<sup>1</sup> | p-value | HR<sup>1</sup> | 95% CI<sup>1</sup> | p-value |
| Eide (2017) | 0.73 | 0.31, 1.74 | 0.5 | 0.69 | 0.28, 1.68 | 0.4 |
| Guinney (2015) | 0.73 | 0.31, 1.74 | 0.5 | 0.69 | 0.28, 1.68 | 0.4 |
| **Mesenchymal** |  |  |  |  |  |  |
| Lehmann (2011) | 0.83 | 0.32, 2.15 | 0.7 | 0.78 | 0.30, 2.07 | 0.6 |
| Phillips (2006) | 0.76 | 0.33, 1.77 | 0.5 | 0.68 | 0.27, 1.73 | 0.4 |
| Walter (2013) | 0.55 | 0.20, 1.47 | 0.2 | 0.51 | 0.18, 1.41 | 0.2 |
| Verhaak (2010) | 0.66 | 0.23, 1.90 | 0.4 | 0.62 | 0.21, 1.81 | 0.4 |
| **Stemness** |  |  |  |  |  |  |
| Ragulan (2019) | 0.77 | 0.39, 1.52 | 0.5 | 0.72 | 0.35, 1.47 | 0.4 |
| Sadanandam (2013) | 0.72 | 0.33, 1.55 | 0.4 | 0.65 | 0.28, 1.49 | 0.3 |
| **Tumor Content** |  |  |  |  |  |  |
| CPE | 4.57 | 0.02, 910 | 0.6 | 4.63 | 0.03, 753 | 0.6 |
| **Stage** |  |  |  |  |  |  |
| I | — | — |  |  |  |  |
| II | 1.16 | 0.36, 3.76 | 0.8 |  |  |  |
| III | 1.09 | 0.29, 4.04 | >0.9 |  |  |  |
| IV |  |  |  |  |  |  |

<sup>1</sup> HR = Hazard Ratio, CI = Confidence Interval

## 3. Discussion

This study examined the impact of the cell type composition of a tumor sample on 11 established gene expression signatures that are proposed to describe stemness, mesenchymal, and EMT cancer subtypes [2–12]. It is noteworthy that not all of the investigated EMT-related signatures have been proposed to serve as readouts of tumor mesenchymality in clinical samples. Previous studies revealed that cell type composition severely affects published transcriptional cancer subtyping procedures with EMT-related characteristics across multiple cancer indications [14, 19, 20, 22, 24]. However, a global examination of the affected signatures and cancer indications was lacking.

We found that across cancer indications, the discussed signatures reveal a strong footprint of epithelial versus mesenchymal characteristics in clinical samples, as indicated by high coherence scores. These high coherence scores were observed not only in the original cancer indications, in which the signatures were developed, but also in other indications. The key question remains as to whether this signal results from differences in tumor characteristics (tumor-intrinsic) or the number of fibroblasts sampled into a tumor specimen (from the tumor microenvironment or from neighboring normal tissue). The tumor content of a clinical tumor sample is strongly associated with high EMT-related signature scores. It became evident that biopsies with high tumor content are less frequently associated with EMT-related phenomena across different cancer tissues than macroenvironment or tumor samples with low tumor content. This finding was corroborated by our analysis of TME-naïve cancer cell lines. The EMT/stemness expression footprint was reduced across cancer indications when stromal or fibroblast cell lines were excluded from the analysis. Although previous studies highlighted the dependence of signature scores on the tumor content of CRC and BRCA samples [15, 20, 37], a comprehensive analysis of cancer indications and established EMT-related signatures was lacking. Our findings indicate that, as a group, these signatures indeed have significant and steady expression footprints; however, they suggest that sampling of fibroblasts from the tumor macro- and microenvironment is the reason for high scores and that tumor-intrinsic mesenchymality can hardly be assessed reliably with these signatures in clinical samples. In other words, the fibroblast-rich cell type composition of a sample with low tumor content might disguise the true cancer phenotype of a clinical tumor sample, eventually affecting the prevalence and characteristics of the other cancer subtypes. Although some papers acknowledged this issue for a subset of the analyzed signatures [14], there is no general recognition. We hypothesize here that this phenomenon could affect expression-based subtyping schemes in many other indications, in which the existence of mesenchymal/stemness phenotypes has been postulated [38, 39].

Consistent with the results in BRCA [20], CRC [19–22], HNC [14], and lung and pancreatic cancer [24], scRNA-seq data indicated that none of the analyzed EMT-related signature scores, and only a minority of signature genes were elevated in tumor cells but rather in fibroblasts. Our integrated RosettaSX analysis of bulk RNA-seq and simulated pseudo bulk samples highlighted that low tumor and fibroblast-enriched pseudo bulk samples had elevated EMT-related and ECM/fibroblast signature scores. In bulk RNA-seq samples, fibroblasts may contribute primarily to the elevated signature scores. Recent studies described leader cells with a hybrid EMT state that can initiate the migration of cancer cell clusters to metastatic sites [40, 41]. This study does not invalidate such a phenomenon, as the number of patients and sampling procedures limited our scRNA-seq analyses. However, it may fail to recognize individual cancer cells with EMT-related phenotypes since a few cancer-specific signature genes cannot guide the signature profile in a complex cell mixture of low tumor-content bulk RNA-seq samples. Our integrated gene expression signature analysis showed that samples with low tumor content closely align with profiles of fibroblast-enriched pseudo-bulk samples, both displaying higher EMT-related signature scores than in any sample with high tumor content. In samples with low tumor content, EMT-related signature scores are driven by the fraction of tumor-surrounding cells, primarily fibroblasts. Thus, accurate control of the tumor content is required to determine cancer-specific EMT-related phenomena using gene expression signatures.

In CRC, research groups reinterpreted the influence of fibroblast content on EMT-related subtypes as a sign of fibrosis [9, 19, 20], rather than an artifact of the sampling procedure. The main argument for fibrotic activity being a tumor trait, and thus an EMT-related subtype, is an association between patient survival and the net signature assignment in survival analyses. We could not verify an association between patient survival and EMT-related signature scores in untreated patients by using TCGA. Using the Pan-Cancer Atlas outcome data [42], our study suggests that the tumor’s degree of invasion into stromal tissue (e.g., tumor stage) drives the association. Therefore, we believe that single observations of differences in survival in stratifications by EMT-related signatures in other studies cannot be regarded as validation of the ability of these signatures to score tumor-intrinsic mesenchymality/stemness characteristics.

## 4. Conclusion

Our findings show that the expression signals of genes and gene expression signatures that are proposed to be associated with EMT or stemness in cancer should be interpreted with caution when they are investigated in clinical tumor samples.

If the experimental procedures are not tightly controlled, the scores of EMT-related signatures in cancer tissues are mostly driven by non-tumor cells such as fibroblasts. Fibroblast content is inversely correlated with tumor content. High fractions of fibroblasts can result from the inclusion of fibroblasts from the tumor microenvironment or from the tumor-adjacent neighboring stroma of the tumor macroenvironment. High scores of EMT-related signatures are often not the result of intrinsic stemness or the mesenchymality phenotype of tumor cells, but rather the result of a sampling artifact.

Consequently, we recommend implementing experimental procedures to maintain high tumor content in clinical gene expression profiling studies with the intent to use EMT-related gene signatures to identify intrinsic tumor stemness or mesenchymality.

## 5. Materials and Methods

### 5.1 Differential gene expression analysis in scRNA-seq data

For the identification of differentially expressed genes the FindMarkers function of the Seurat package was used. We filtered for genes that had an average fold-change of 2, and a p-value below10*e*^-10^ for a cell type of interest and all other available cells.

### 5.2 Definition of gene expression signatures for the CMS4 CRC subtype

Guinney et al. and Eide et al. implemented two independent models (CMSclassifier and CMScaller) to classify consensus molecular subtypes (CMS; the CMS4 subtype is associated with EMT) [9, 12]. However, there was no signature provided for these subtypes, and consequently, we selected gene sets based on differential gene expression analysis of TCGA CRC bulk sequencing data. For the CMSclassifier, we downloaded the CMS sample labels from Synapse [49], and for CMScaller, we used the R package for the prediction of CMS subtypes on our expression data. Samples that could not be classified into any CMS subtype were excluded from analysis. Genes differentially expressed in CMS4 samples compared to all other samples were used as a gene expression signature with a maximum of 100 genes, a p-value <= .01, and a log2 fold change larger than 2.

### 5.3 Definition and scoring of gene expression signatures

Count data were normalized using the trimmed mean M-values (TMM) normalization of the edgeR [44] R-package using default parameters. Gene expression signatures were studied using methods recently published in the context of our RosettaSX platform for expression signature investigation [25].

To calculate the signature scores in the scRNA-seq analyses, we used the Seurat addModule function. For signature scores of bulk RNA-seq data, we used the mean TPM of TMM normalized expression, to describe the activity of a signature. For the association statistics, we used Pearson’s correlation. We used Euclidean distance and complete hierarchical clustering to visualize the heatmaps.

### 5.4 Judging the coherence and translatability of signatures by the coherence score

To determine the relevance of the signatures for an expression dataset, we applied the coherence score (CS) [25, 26], and the average Pearson correlation coefficient of all signature gene pairs in a specific dataset. This score indicates a strong negative or positive coherence (CS close to 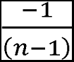 or +1) between all gene expression profiles of a signature or no linear relationship (CS=0).

### 5.5 Definition of pseudo bulk sequencing data

To create pseudo-bulk RNA-seq data from scRNA-seq data, we the SimBu R package [50] and generated pseudo-bulks comprising fibroblast and malignant cells. We complied with previously described methods to integrate pseudo bulk and TCG bulk samples [51]. Briefly, we normalized bulk RNA counts for gene length and used TMM correction in edgeR (version 3.36.0) for both pseudo bulk and bulk samples.

### 5.6 Survival analysis

Outcome data were downloaded from the Pan Cancer Atlas [42]. Due to the recommendation of the Pan Cancer Atlas study, we could only analyze patients with COAD and BRCA. Univariate Cox proportional hazards models were individually fitted to each signature. Multivariate models were fitted using AJCC staging as covariates. We tested for proportional hazards using the Schoenfeld residuals.

### 5.7 Data Availability

Gene expression RNA-seq data were downloaded from the Xena database [43]. CPE tumor purity scores were assessed using TCGAbiolinks [45]. Cancer cell line data were downloaded from DepMap [46] (release 20q4). For cell line annotations, we used integrated Cellosaurus cell line annotations [47].

We downloaded scRNA-seq data (EMTAB8107, GSE148673, GSE161529, EMTAB8107, GSE146771, GSE166555, GSE141383, and GSE103322) from the TISCH2 database [48]. Each dataset was analyzed independently, and the cell annotations of the original authors were used.

## 6 Acknowledgement

The authors thank Michail Yekelchyk and Sven-Eric Schelhorn for providing access to the TISCH2 and TCGA data.

## 8. Supplementary Material

**Supplementary Figure S1:**
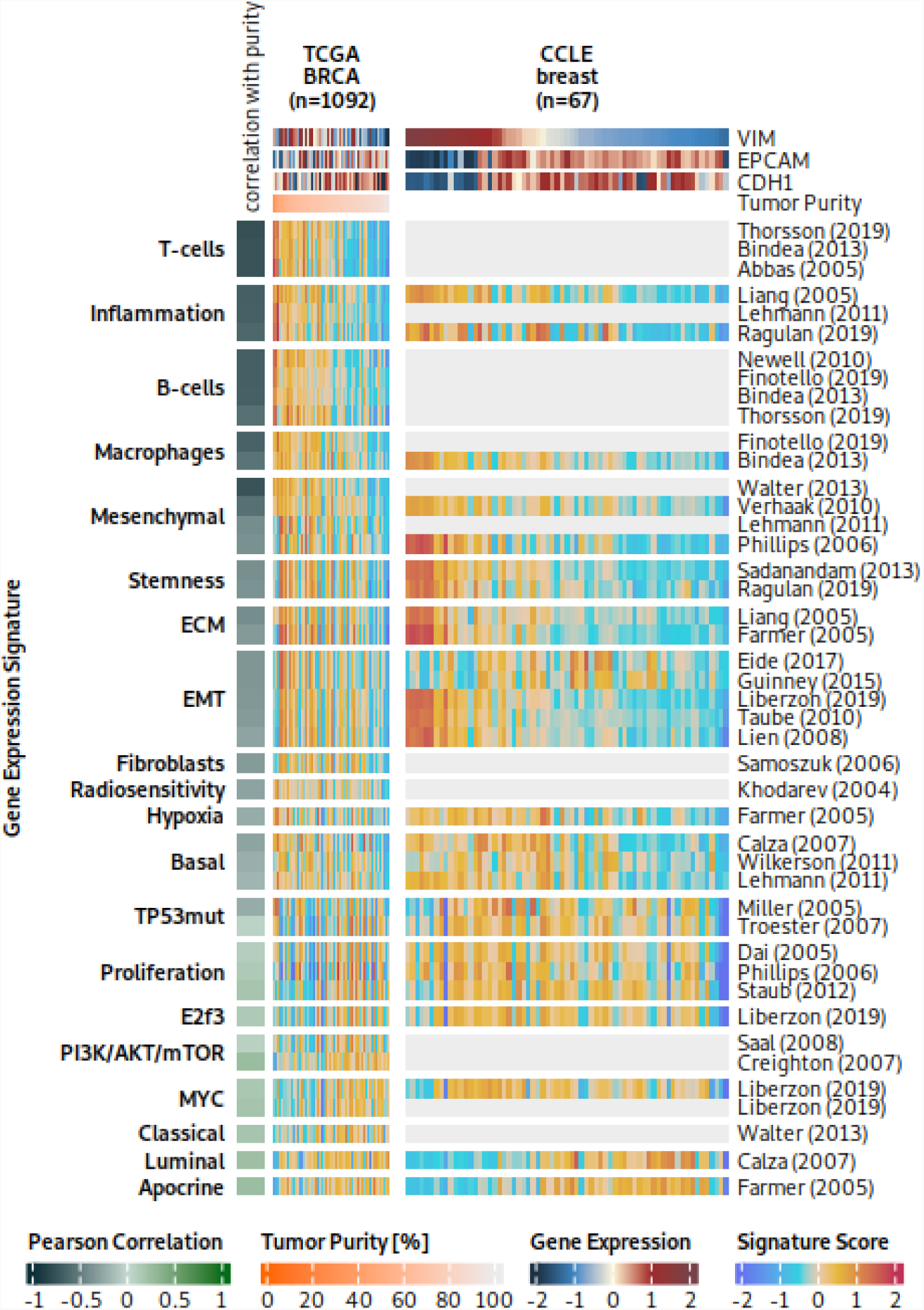
RosettaSX analysis for forty-four filtered gene expression signatures in breast TCGA RNA-seq samples and DepMap breast cancer cell lines. Cell lines with elevated EMT-related signature scores have a mesenchymal origin (HS281T, HS343T, HS606T, HS578T, HS274T)

**Supplementary Figure S2:**
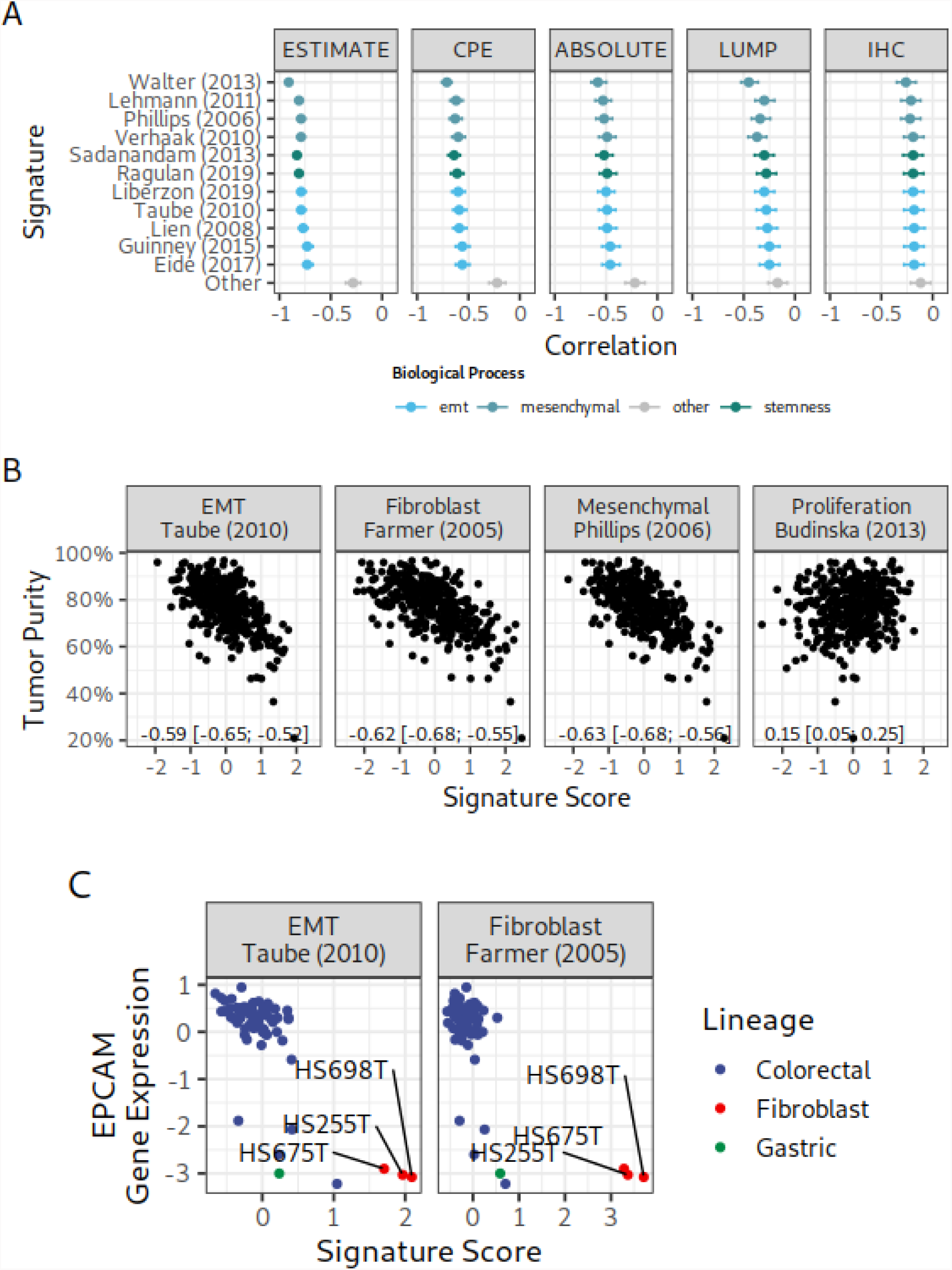
Comparison of signature scores with tumor content in TCGA CRC and epithelial markers in TME-naïve colorectal cell lines A: Association between gene expression scores for mesenchymal, EMT, stemness signatures, and tumor purity in TCGA CRC. The ‘others’ category summarizes the average correlation, lower and upper 95th percentile of thirty-three coherent gene expression signatures. Colors indicate different biological processes, and error bars indicate the upper and lower 95th percentile confidence intervals. B: Scatter plots comparing mesenchymal, stemness, EMT, and cancer-related gene expression signatures with tumor purity. C: Scatter plot comparing an epithelial marker (EPCAM) with EMT and fibroblast signature scores in CRC cell lines, colored by cell line lineage.

**Supplementary Figure S3:**
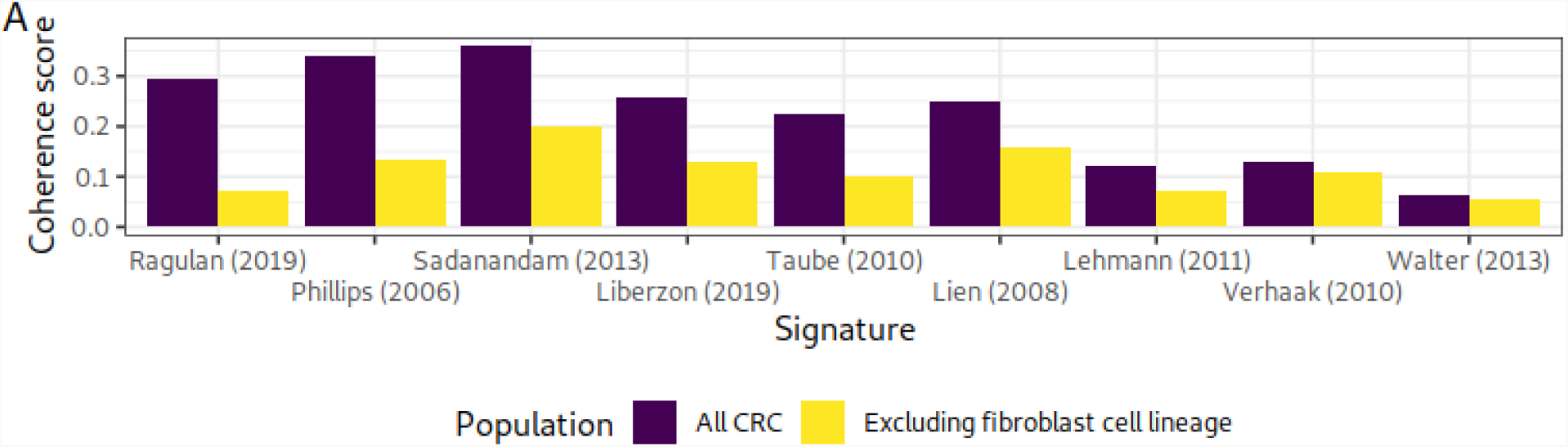
Coherence score for gene expression signatures in colorectal cancer cell lines strongly affected by mesenchymal origin lines. While red bars represent the correlation coefficient for all seventy-four cell lines, the blue bars show the coefficient for the 71 cell lines (excluding lines with fibroblast lineage).

**Supplementary Table S1:**
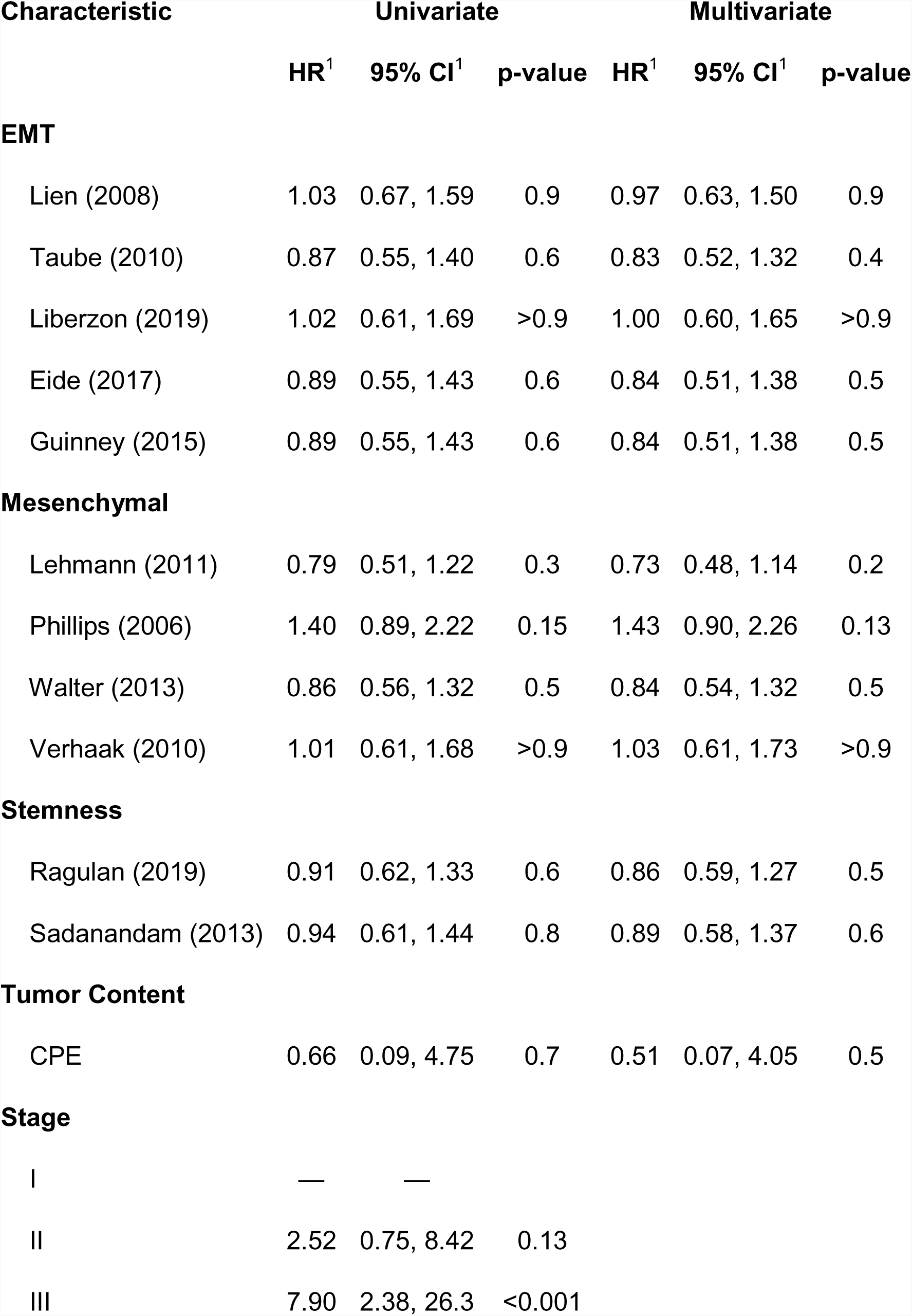
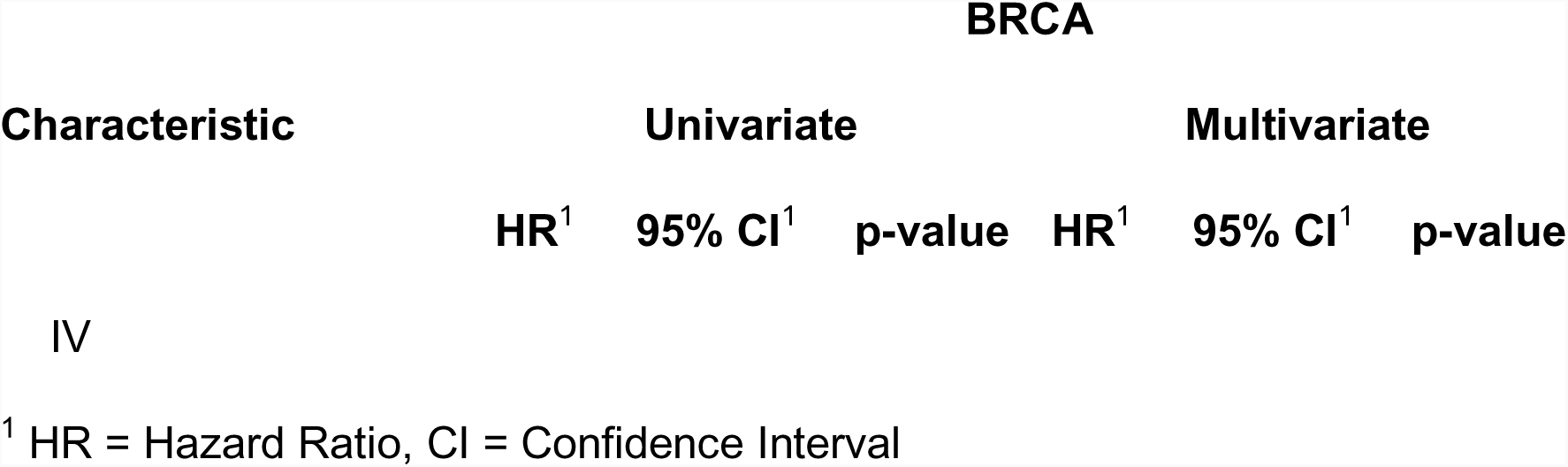
Cox proportional hazards analysis for EMT-related signature scores and TCGA cancer patients. We show the results of univariate and multivariate Cox regression models to analyze the influence of EMT-related gene expression signature scores and stage annotation on progression-free interval survival of TCGA BRCA patients. Multivariate analysis was performed using tumor stage as a covariate.

BRCA
| Characteristic | Univariate |  |  | Multivariate |  |  |
| --- | --- | --- | --- | --- | --- | --- |
|  | HR<sup>1</sup> | 95% CI<sup>1</sup> | p-value | HR<sup>1</sup> | 95% CI<sup>1</sup> | p-value |
| **EMT** |  |  |  |  |  |  |
| Lien (2008) | 1.03 | 0.67, 1.59 | 0.9 | 0.97 | 0.63, 1.50 | 0.9 |
| Taube (2010) | 0.87 | 0.55, 1.40 | 0.6 | 0.83 | 0.52, 1.32 | 0.4 |
| Liberzon (2019) | 1.02 | 0.61, 1.69 | >0.9 | 1.00 | 0.60, 1.65 | >0.9 |
| Eide (2017) | 0.89 | 0.55, 1.43 | 0.6 | 0.84 | 0.51, 1.38 | 0.5 |
| Guinney (2015) | 0.89 | 0.55, 1.43 | 0.6 | 0.84 | 0.51, 1.38 | 0.5 |
| **Mesenchymal** |  |  |  |  |  |  |
| Lehmann (2011) | 0.79 | 0.51, 1.22 | 0.3 | 0.73 | 0.48, 1.14 | 0.2 |
| Phillips (2006) | 1.40 | 0.89, 2.22 | 0.15 | 1.43 | 0.90, 2.26 | 0.13 |
| Walter (2013) | 0.86 | 0.56, 1.32 | 0.5 | 0.84 | 0.54, 1.32 | 0.5 |
| Verhaak (2010) | 1.01 | 0.61, 1.68 | >0.9 | 1.03 | 0.61, 1.73 | >0.9 |
| **Stemness** |  |  |  |  |  |  |
| Ragulan (2019) | 0.91 | 0.62, 1.33 | 0.6 | 0.86 | 0.59, 1.27 | 0.5 |
| Sadanandam (2013) | 0.94 | 0.61, 1.44 | 0.8 | 0.89 | 0.58, 1.37 | 0.6 |
| **Tumor Content** |  |  |  |  |  |  |
| CPE | 0.66 | 0.09, 4.75 | 0.7 | 0.51 | 0.07, 4.05 | 0.5 |
| **Stage** |  |  |  |  |  |  |
| I | — | — |  |  |  |  |
| II | 2.52 | 0.75, 8.42 | 0.13 |  |  |  |
| III | 7.90 | 2.38, 26.3 | <0.001 |  |  |  |

| BRCA |  |  |  |  |  |  |
| --- | --- | --- | --- | --- | --- | --- |
| Characteristic | Univariate |  |  | Multivariate |  |  |
|  | HR<sup>1</sup> | 95% CI<sup>1</sup> | p-value | HR<sup>1</sup> | 95% CI<sup>1</sup> | p-value |

**Supplementary Figure S4:**
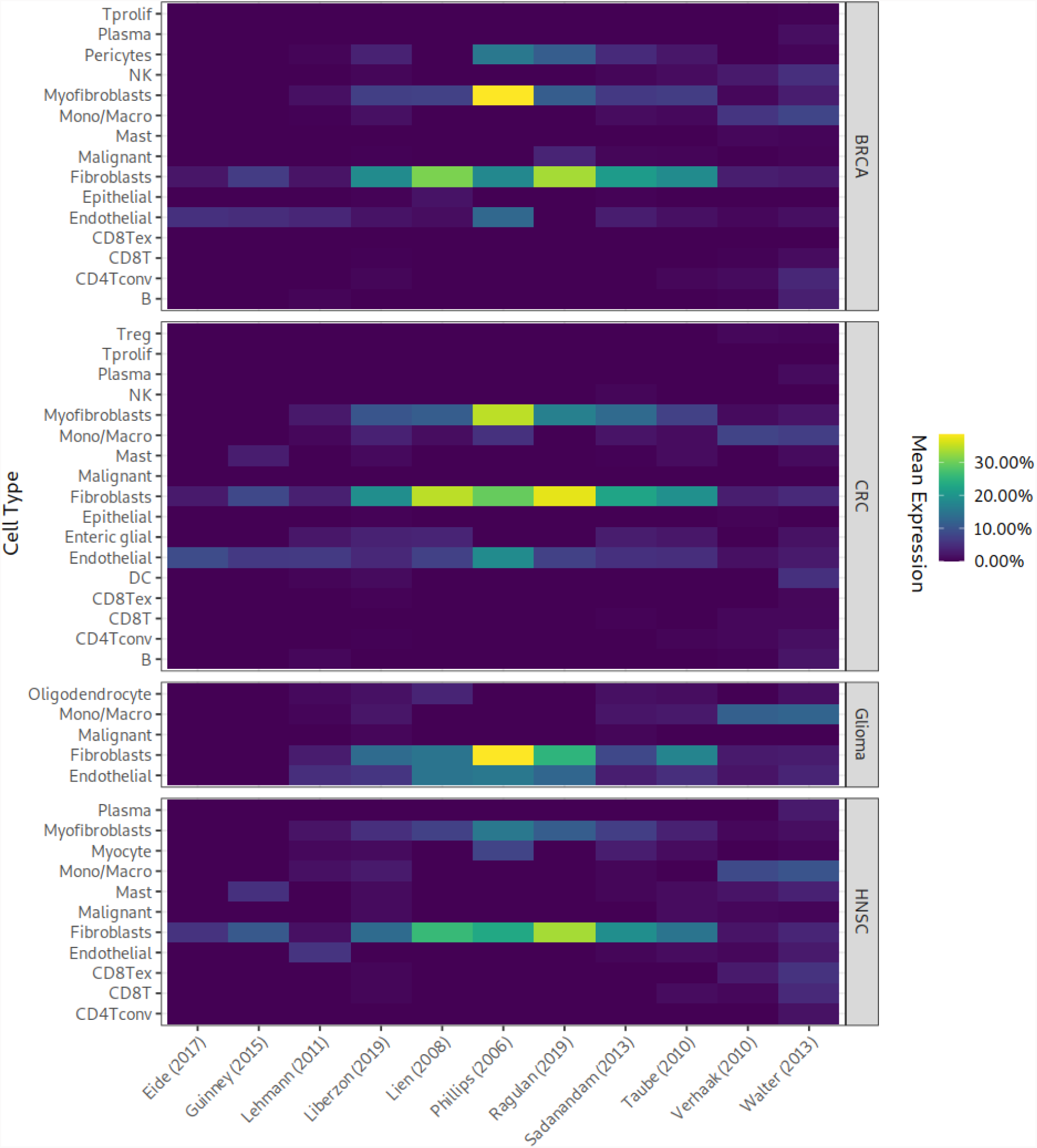
Percentage of differentially expressed genes in BRCA, CRC, HNSC, and Glioma. Each cell-type was analyzed in comparison to all other cell lines within the respective dataset. Percentages were averaged across datasets for the same indication.

